# Versatile proteome labelling in fruit flies with SILAF

**DOI:** 10.1101/590315

**Authors:** Florian A. Schober, Ilian Atanassov, David Moore, Anna Wedell, Christoph Freyer, Anna Wredenberg

## Abstract

*Drosophila melanogaster* has been a working horse of genetics and cell biology for more than a century. However, proteomic-based methods have been limited due to technical obstacles, especially the lack of reliable labelling methods. Here, we advanced a chemically defined food source into stable-isotope labelling of amino acids in flies (SILAF). It allows for the rapid generation of a large number of flies with full incorporation of lysine-6. SILAF followed by fractionation and enrichment gave proteomic insights at a depth of 5,966 proteins and 7,496 phosphorylation sites, which substantiated metabolic regulation on enzymatic level. Furthermore, the label can be traced and predicts protein turnover rates during different developmental stages. The ease and versatility of the method actuates the fruit fly as an appealing model in proteomic and post-translational modification studies and it enlarges potential metabolic applications based on heavy amino acid diets.

## INTRODUCTION

Stable-isotope labelling of amino acids in cell culture, known as SILAC, uses non-radioactive isotopic labelling to detect differences in relative abundance between at least two peptide populations simultaneously (1). This allows accurate peptide and protein quantification even upon fractionation of peptide mixes and thus greatly enhances the depth of large-scale screening for proteins, post-translational modifications (2) or protein-protein interactions (3, 4). The simplicity and robustness of SILAC has led to its broad adoption in a variety of biological specialities and has been applied to a number of animal models (5, 6). In *Drosophila melanogaster* (Dm) SILAC labelling was originally reported in 2010 by using labelled yeast as food source (7). Although technically feasible, it has not been adopted by the fly community, due to the high amounts and costs of stable isotopes required, labelling efficiencies at the borderline of SILAC usability and a general inflexibility concerning the usage of several different heavy amino acids. A follow-up study in 2013 (8) could decrease the labelled yeast mass profoundly, whilst retaining 95% labelling efficiency. Yet, the authors describe an undesirable conversion of amino acids isotopes that caused a decreased peptide identification rate. Although normalisation on a protein level can account for this, studies relying on accurate quantification of peptides, e.g. post-translational modification scans have not been feasible in *Drosophila* yet. Furthermore, the somewhat unpredictable yeast metabolism does not allow precise tracing of amino acid labels in order to reliably predict protein turnover rates and complicates the introduction of several labels or additive substrates at the same time. Here, we present an alternative, direct labelling approach by employing a fully defined chemical food source that contains heavy amino acids isotopes, which addresses several key concerns of stable-isotope labelling in fruit flies.

## RESULTS

To overcome the difficulties associated with traditional yeast-based food in SILAC, we used a holidic food source in this study containing 47 ingredients (9, 10), including defined amounts of eight essential amino acids. Flies grown on this holidic food are viable and fertile with a comparable lifespan, although they do present with a prolonged developmental larvae stage, pupating after approximately nine instead of five days after egg laying (dae) at 25°C (9, 10).

### SILAF enables full amino acid labelling

In its more popular application in cell culture, SILAC relies on labelling with both “heavy” ^13^C_6_-lysine and ^13^C_6_^15^N_4_-arginine followed by digestion with trypsin, which cuts C-terminally of both amino acids. However, previous results in both cells and the fly suggested that arginine can be metabolised into other amino acids (8, 11). In agreement with this, growing larvae on holidic food containing heavy-lysine and -arginine, followed by protein identification using liquid chromatography-tandem mass spectrometry (LC-MS/MS), identified a number of heavy [^13^C_5_^15^N_1_] proline-containing peptides (Supplementary Fig. 1a, b). In addition, we observed a decrease in the average intensity of the heavy fraction in a peptide mixture from heavy and light holidic food indicating a “loss” of the heavy label (Supplementary Fig. 1c). We therefore switched to a holidic medium containing only heavy-lysine and performed protein digestion with the endoproteinase Lys-C. This equalised heavy and light intensities in a control experiment (Supplementary Fig. 1c), indicating the absence of metabolic turnover and incorporation into other amino acids. Reassuringly, the usage of Lys-C as opposed to trypsin did not compromise peptide identification (Supplementary Fig. 1d).

In order for SILAF to be used in a proteomic experiment, it is desirable to achieve full incorporation of the heavy amino acid in the proteome and labelling rates above 95% are required to perform quantitative proteomic experiments (12). To measure incorporation efficiency of heavy-lysine, we allowed wild type *white Dahomey* (wDah) flies, which had been grown on standard yeast food, to lay eggs for eight hours on holidic food containing heavy-lysine. Labelling efficiency was assessed during Dm development in two-day intervals by LC-MS/MS. Heavy-lysine was incorporated rapidly and already at four days after egg-lay (dae) we reached 97.2% incorporation efficiency. By six dae we reached 99.5% incorporation (Fig. 1a), which is above the stated purity of the heavy amino acid of 99%, and we therefore achieved full labelling of larvae and flies. In contrast to previous methods using labelled yeast (13), holidic food labelling makes it possible to also confidently use second and third instar larva stages as well.

**Figure 1:**
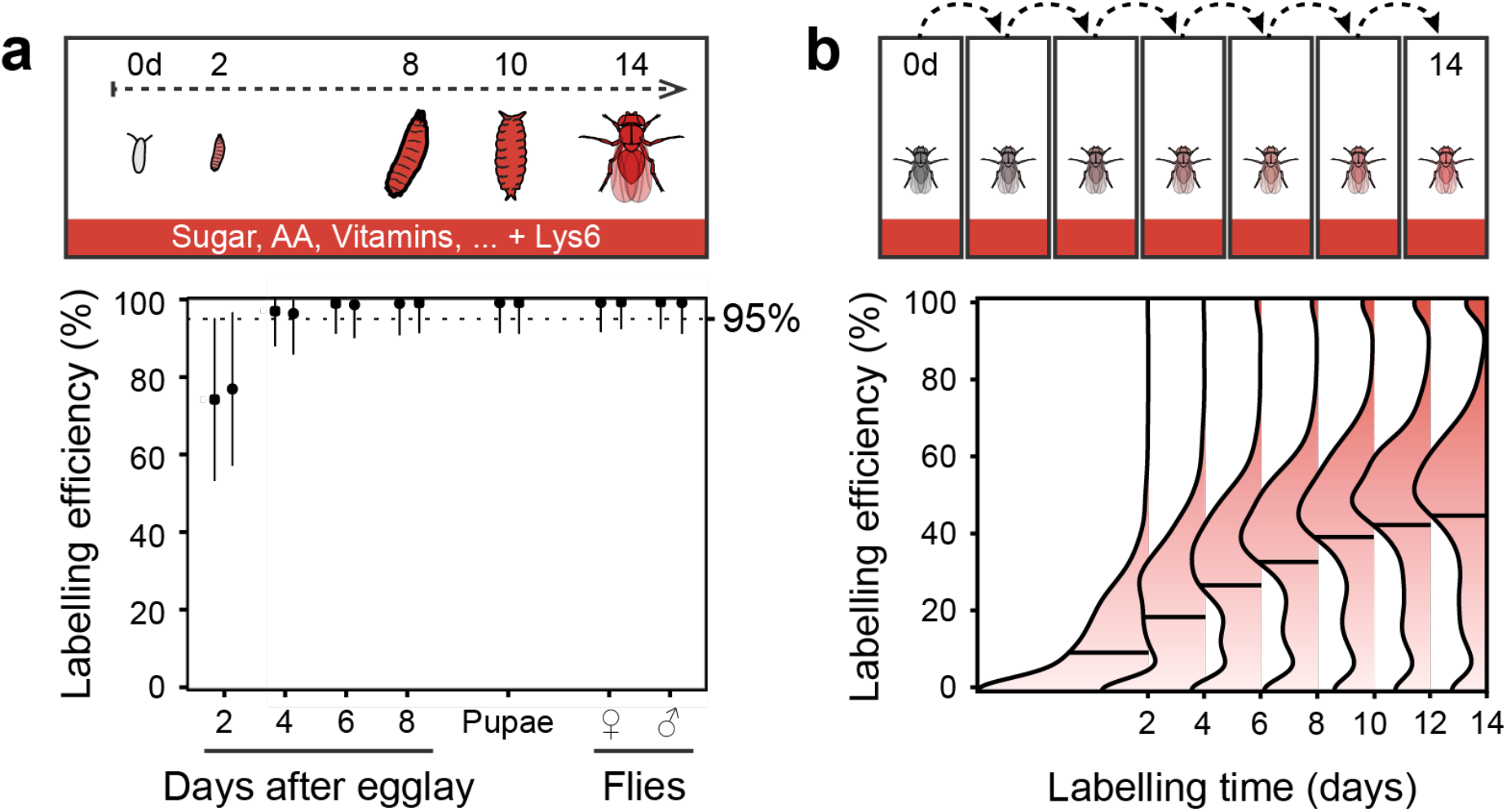
SILAF labelling in developing and adult flies. (a) Protein incorporation rates of Lys-6 in developing flies were measured every other day starting from unlabelled fertilised eggs to one day-old flies. Dotted line indicates 95% labelling efficiency. Mean ± s.d. are shown for two replicates per time point. (b) Density curves of Lys-6 incorporation rates in adult flies were measured from a one-day old unlabelled fly, for 14 days, in 2-day intervals. Black lines are mean values of two replicates.

Once we established the complete incorporation of heavy-lysine in larvae from which we obtained fully labelled adult flies, we tested the utility of holidic food-based SILAF for quantitative proteomics. We first set out to evaluate the precision of quantification and compared it to label-free quantification (LFQ). We obtained flies from larvae grown on either light or heavy holidic food, mixed the protein extracts and performed single-shot LC-MS/MS analysis. We observed a high average (>0.98) Pearson’s correlation coefficient of the intensities of the heavy and light proteins (Supplementary Fig. 1f), indicating high precision of the technique, low or no conversion of heavy lysine into other amino acids and absence of artefacts related to labelling. Interestingly, we achieved similar correlation coefficient values and average standard deviation between replicates by LFQ on the light fraction of these samples (Supplementary Fig. 1e,g), indicative of the advancement of modern single-shot label-free analysis.

### Lysine-6 incorporation is a qualitative readout for protein turnover

Next, we evaluated the labelling efficiency in adult flies. We transferred hatched wild type wDah flies grown on standard yeast food to heavy-lysine containing holidic food and collected samples in two-day intervals for LS-MS/MS analysis. Compared to larvae, adult flies showed a much lower rate of heavy label incorporation, reaching a maximum of 47.0% after 14 days (Fig. 1b). The distinction likely originates from differences in the metabolic needs of the two developmental stages. Larvae are subjected to constant growth and the rapid heavy label incorporation reflects a dilution of the original light amino acid pool, while the adult fly consists of primarily post mitotic tissue, representing a system with balanced rates of synthesis and degradation, leading to a slow and steady heavy label incorporation. In this line, testing the incorporation of the heavy label in the adult fly effectively represents the pulse phase of a pulse-chase experiment, indicating that SILAF can be used to determine protein turn-over rates *in vivo.* Lysine-6 incorporation occurs in a non-uni-form manner (Fig. 1b) and we therefore examined the proteome wide incorporation of the heavy label in flies at labelling day 14 (Fig. 2a). Overrepresentation analysis (ORA) against KEGG metabolic maps (14) identified low levels of labelling for proteins involved in oxidative phosphorylation, whereas ribosomal protein incorporation rates were close to 50%.

**Figure 2:**
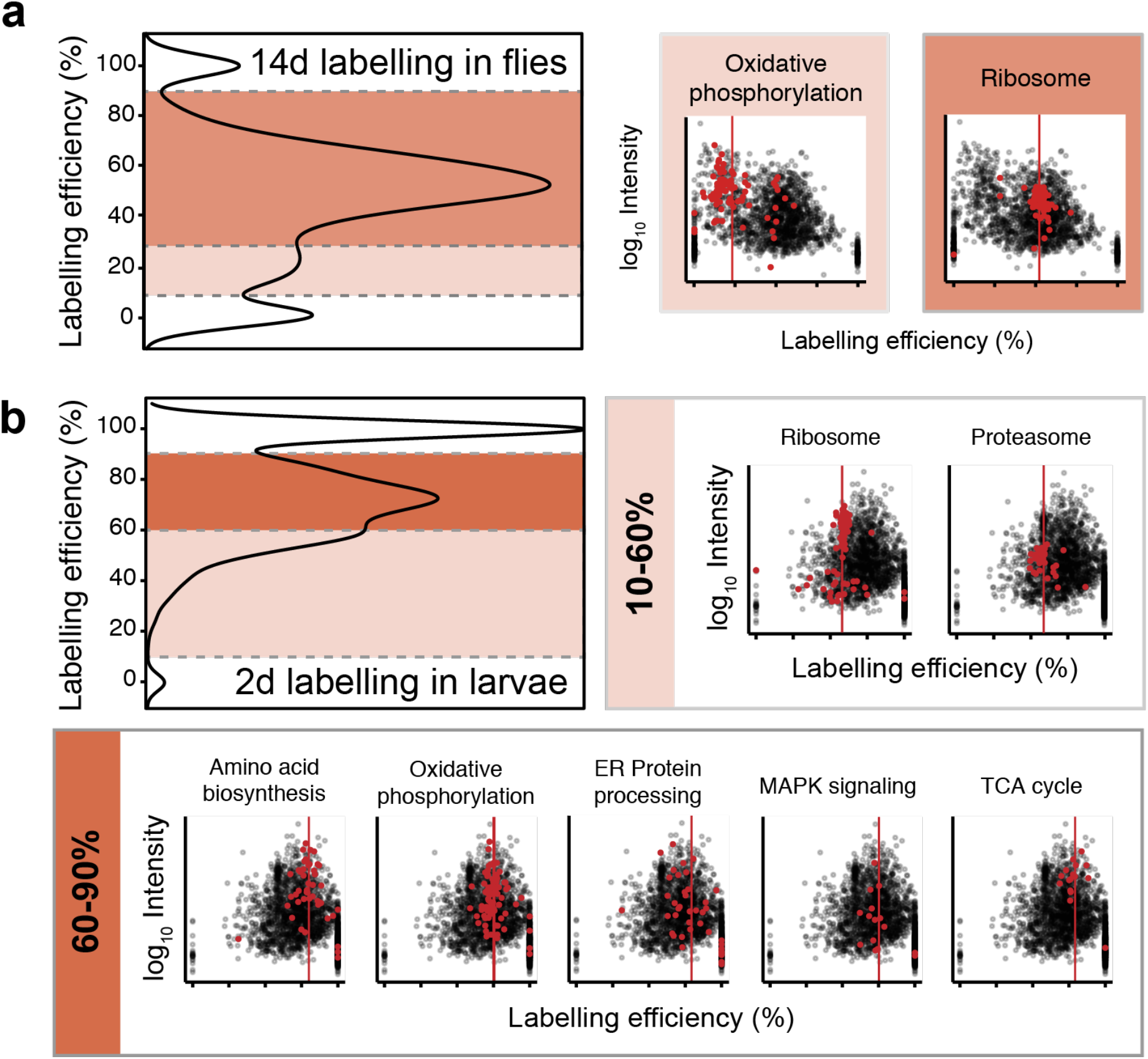
Lys-6 tracing in developing larvae and adult flies. (a) Unlabelled flies were put on heavy holidic food. Labelling profile after 14 days of labelling as a mean of two replicates with the identified fractions, of which significantly enriched KEGG pathways derived by overrepresentation analysis are shown. Labelling efficiency was divided into three bins: <10% or >90% labelling of peptides (white), 10-20% (light red) and 30-90% (dark red). KEGG pathway overrepresentation analysis was done on proteins in the 10-20% and 30-90% bins. Proteins belonging to significantly enriched KEGG terms are highlighted (red) with mean value as the red line on labelling efficiency vs protein MS intensity plots of one replicate. (b) Density curve showing Lys-6 incorporation into proteins in larvae 2 dae (top left). Incorporation values were divided into three bins: <10% or >90% (white), 10-60% (light red) and 60-90% (dark red). Analysis on each bin and depiction of significant pathways as in (a).

Similar to the analysis of adult flies, we observed a non-uniform distribution in the incorporation density curve at two dae (Fig. 2b). In contrast to adult labelling, we found components of energy conversion pathways, i.e. oxidative phosphorylation and tricarboxylic acid cycle (TCA) in a prominent peak between 60-90% labelling efficiency, whereas members of the protein turnover machinery, i.e. ribosome and proteasome clustered lower at around 50% labelling efficiency. The different heavy label incorporation rates in larvae and flies therefore seem to reflect differences in synthesis rates and/or initial amount of the respective protein groups *in vivo.*

### Small-scale fractionation with SILAF gives deep proteomic insights

To assess SILAF for its biological application we obtained male and female flies from larvae, grown on either heavy or light holidic food, mixed the protein extracts and performed LC-MS/MS analysis. In order to achieve more comprehensive proteome coverage, we performed small-scale high pH reversed phase peptide fractionation on 20μg of input peptides. The direct labelling-based comparison of male and female flies detected 4,577 proteins and revealed profound differences in the abundance of a number of sex related proteins (Supplementary Fig. 2a), including yolk sack proteins, which are exclusively expressed in the female fat body (15). A gene set enrichment analysis against gene ontologies revealed functional categories related to sex differentiation and mating (Supplementary Fig. 2b). We further compared our results with those from Sury and colleagues (7) and observed good agreement in the level of regulation of sex related proteins (Supplementary Fig. 2c). This indicates that experiments based on holidic food give comparable results to traditional yeast-based labelling, while at the same time circumventing previous technical obstacles.

### Single-shot SILAF proteome analysis can be used to identify the aetiology of genetic defects

We applied SILAF to study the proteomic alterations in a pathological fly model. We chose flies with silenced expression of the leucine rich pentatricopeptide repeat containing protein DmLRPPRC1, also known as the bicoid-stability factor (BSF); a mitochondrial protein involved in mitochondrial transcript stability and polyadenylation (16, 17). Mutations in human *LRPPRC* have been associated with mitochondrial disease (18, 19), and knockdown of DmLRPPRC1 by RNAi in flies (DmLRPPRC1^KD^, Fig. 3a) leads to a severe mitochondrial respiratory chain deficiency, delayed larval development and decreased life span (16).

**Figure 3:**
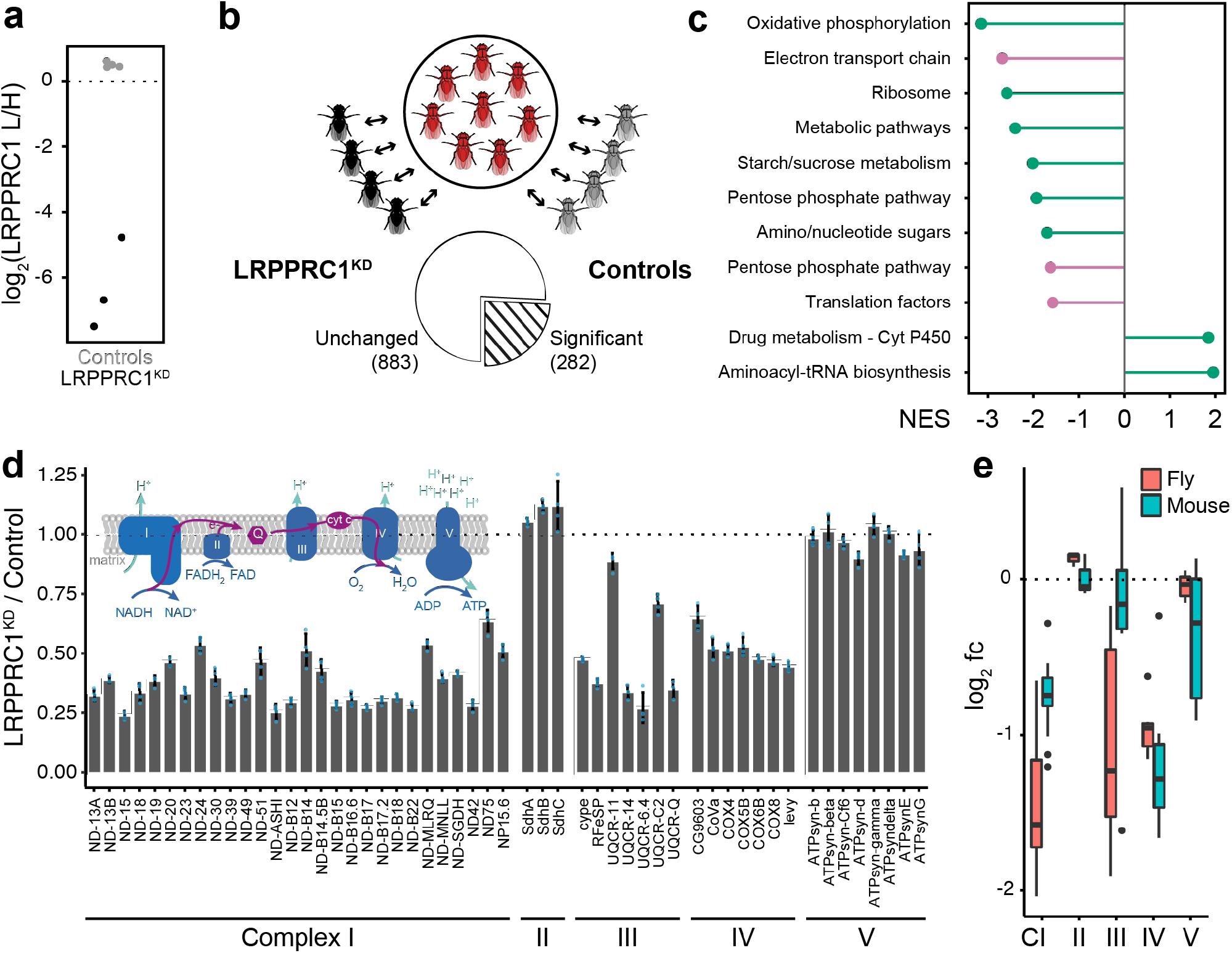
SILAF application to quantify proteomic changes in a DmLRPPRC1 knock-down model. (a) Protein intensities of DmLRPPRC1 determined on newly hatched flies. One replicate without determined protein levels. (b) SILAF spike-in based on a common heavy standard and ratio of ratios comparison. Protein extracts from light adult control and knock-down flies, n=4, both raised on yeast diet, were each mixed with an equal amount of pooled protein extracts from flies grown on a heavy SILAF diet. Protein mixtures were analysed by single-shot LC-MS/MS, followed by a two-sample moderated t-test. Significance threshold was set at an FDR of 0.1%. (c) Gene Set Enrichment analysis of proteins shown in Fig. 1c against KEGG (green) and Wikipathway (pink) reference database. (d) Protein levels of OXPHOS complexes I (NADH:ubiquinone oxidoreductase), complex III (ubiquione:cytochrome c oxidoreductase), and complex IV (cytochrome c oxidase) were significantly reduced, while subunits of complex II (succinate:ubiquinone oxidoreductase) and complex V (ATP synthase) were not affected (SILAF spike-in experiment). Mean ± s.d. and corresponding dotplot (blue) are shown for four replicates. (e) Comparison of OXPHOS complex I to V protein levels obtained by ratio/ratio SILAF in this study and perviously published data obtained by label-free quantification on mitochondrial fractions of 7-week old LRPPRC knock-out mice (21).

We performed single-shot LC-MS/MS analysis on control and DmLRPPRC^KD^ third instar larvae, grown on heavy or light holidic food, respectively (Supplementary Fig. 3a). Alternatively, we adapted a spike-in method (20), where a common protein sample from flies grown on heavy holidic food is spiked in the protein samples from one-day-old flies grown under standard growth (yeast food) conditions. Control and DmLRPPRC1^KD^ flies grown on yeast diet were picked one day after hatching, protein samples were spiked-in with reference protein extracts from a pool of wDah flies grown on heavy holidic medium and subjected to single-shot LC-MS/MS analysis (Fig. 3b). Differential protein expression analysis was performed by comparing the differences to the heavy pool across all samples. Comparison of both approaches showed good correlation (Supplementary Fig. 3b), identifying approximately 1,200 quantified proteins, whereof 76 (6.1 %) or 282 (24.2%) were significantly regulated at 0.1% false discovery rate (FDR) in the classic or spike-in experiments, respectively (Fig. 3b and Supplementary Fig. 3a). A gene set enrichment analysis identified the KEGG pathway “oxidative phosphorylation” as the most negative and significant category (Fig. 3c and Supplementary Fig. 3c,d). Splitting up the pathway into complexes of the electron transport chain and the ATPase showed that only complex I, III and IV subunits were profoundly changed (Fig. 3d), which is in agreement with previous work on this model (16), as well as with a published proteomic dataset on a mouse model of LRPPRC (21) (Fig. 3e).

### Quantification of the proteome and phospho-proteome using SILAF predicts perturbations in metabolic pathways

As part of its design, SILAF allows for samples of interest to be combined and processed together very early on in a proteomic experiment. This makes the approach ideal for quantitative analyses of protein post-translational modifications *in vivo,* especially when combined with some form of enrichment and/or fractionation. We employed large-scale high pH reversed phase fractionation (22) and TiO2 based phospho-peptide enrichment (23) to perform quantitative proteome and phosphoproteome analysis on one-day old flies grown either on standard yeast food or on holidic food (Fig. 4a). In a replicated experiment we quantified up to 4,749 proteins and detected 7,496 phospho-sites (Fig. 4b,c, Supplementary Fig. 4a), with the vast majority modified on serine residues (Supplementary Fig. 4c). Differential expression analysis of the fractionated proteome revealed 141 proteins significantly regulated on the protein level at an FDR of 0.1% (Fig. 4b), corresponding to 3.6% of the total dataset. Among those, the phosphoenolpyruvate carboxy kinase (PEPCK), the rate-limiting enzyme in gluconeogenesis, was increased 3.7-fold on protein level in flies grown on holidic food (Fig. 4b), while components of glycolysis and the oxidative phosphorylation pathway were decreased. Additionally, fatty acid metabolic enzymes were down-regulated as opposed to up-regulated ribosomal and proteasomal components (Fig. 3b, Supplementary Fig. 4d). Combining the phospho-proteome with peptide abundances allowed us to quantify the occupancy of 1,356 phospho-sites from 629 proteins (Supplementary Fig. 4b). Proteins with up-regulated occupancy included factors involved in glucose metabolism with myofibrils as an overrepresented category, whereas components around proteasome and ribosome showed reduced phosphorylation occupancy (Fig. 4c and Supplementary Fig. 4e,f). These findings together point towards a decreased usage of glucose in glycolysis, decreased fatty acid metabolism and an increased turnover and energetic usage of amino acids due to the holidic food, feeding gluconeogenesis (Fig. 4d).

**Figure 4:**
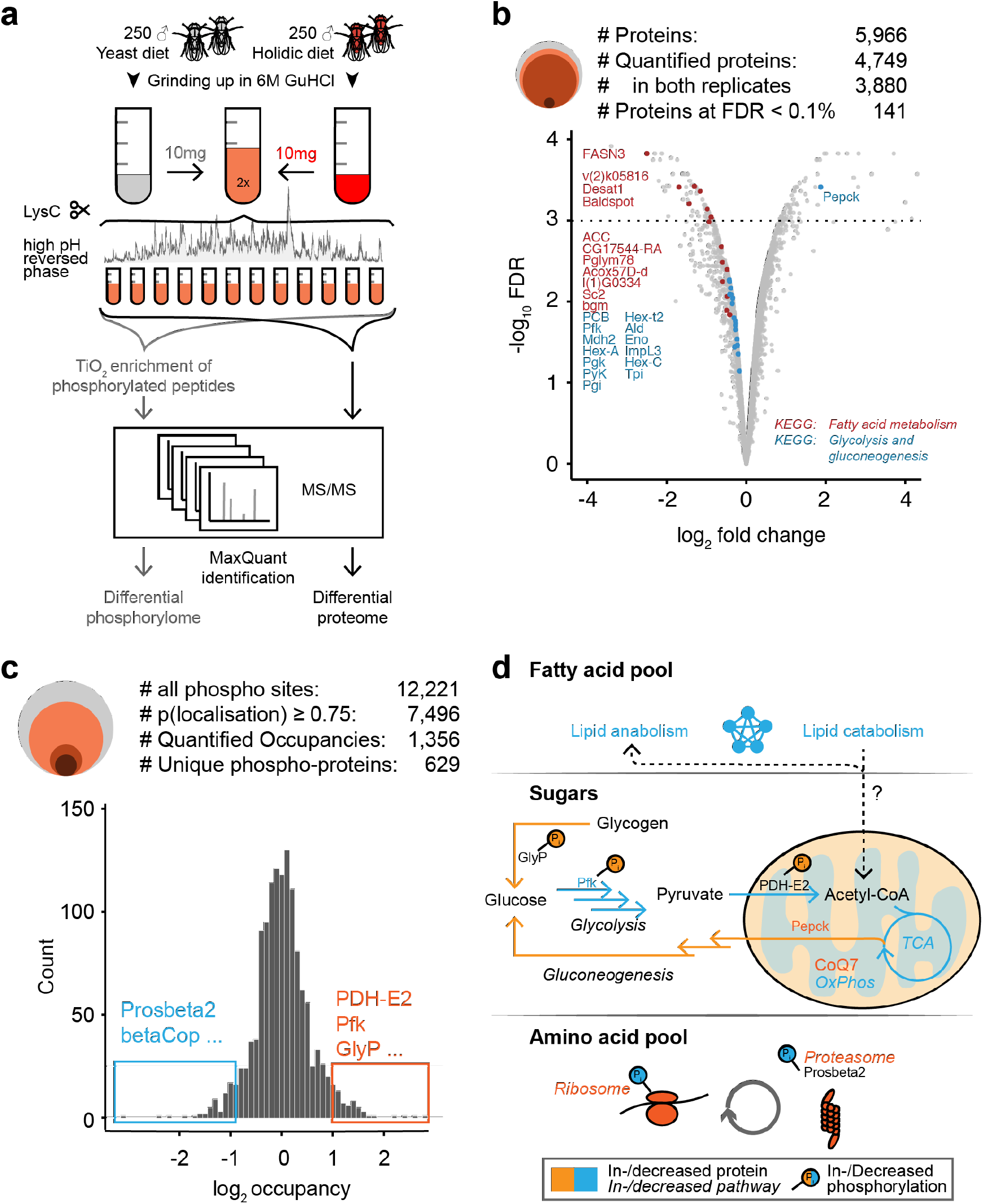
SILAF application to quantify proteome and phosphorylome differences between holidic versus yeast food-grown flies. (a) Proteomic work-flow to obtain a deep proteome and phosphorylome dataset. Protein extracts, from flies grown on light yeast diet or heavy holidic diet, n=2, were mixed, digested with endoproteinase LysC and separated by high pH reversed phase chromatography. Fractionated peptides were either analysed directly by LC-MS/MS (proteome) or subjected to titanium dioxide enrichment (phosphorylome). (b) Total number of identified proteins (grey), proteins quantified in both replicates (orange, and significant proteins (FDR threshold 0.1%, light red) are shown (upper panel). Volcano plot of the proteome comparison with members of two KEGG pathways highlighted. Dotted line indicates the FDR 0.1% threshold. (c) Total number of identified phospho-sites (grey), phospo-sites with high localization probability (orange), phospho-sites with reported occupancy (light red) and number of unique proteins with phospho-site occupancy (dark red) are shown (upper panel). Significant increased (red) or decrease (blue) in phosphor-site occupancy (lower panel). (d) Proposed model of metabolic differences in *Drosophila* on a holidic food diet compared to a standard yeast diet.

## DISCUSSION

Label-free quantification by LC-MS/MS has made immense advances in recent years in accuracy and reproducibility, making many previously impossible analytical investigations now feasible (24). Additionally, it has led to a reduction in the dependency on techniques, such as SILAC, as a quantitative view of the proteome can now also be obtained by LFQ. In fact, we find that SILAF is only marginally more reproducible than LFQ comparisons. However, for many applications that include fractionation or tracing, metabolic labelling with amino acid isotopes is inevitable and the most accurate gold-standard to date (25). SILAC in flies has been evaluated in several reports (7, 8, 13), which were all based on feeding flies with auxotrophic, Lys-8 labelled yeast. In principle, this allowed following a fly lab’s standard experimental routines, replacing normal with labelled yeast in a default medium. However, the amounts of yeast are difficult to obtain both technically and financially (8) and consequently experimental procedures were changed either towards feeding yeast paste or dried yeast (7, 13) or reduced yeast content in solid medium (8). Despite remaining costly (26), labelling ranged from 93.7% to 97.7% and it had been shown that the reason for not achieving complete labelling has been yeast and not fly metabolism (7). Furthermore, as yeast had to be labelled first, comparable to *E. coli* labelling in *C. elegans*-SILAC (27), amino acids are exposed to potential metabolic modifications in yeast cells. In fact, Chang *et al.* observed undesirable conversion of both lysine and arginine into other amino acids in the fly SILAC proteome that negatively affects accurate quantification on peptide level (8).

Here, we present a method that takes advantage of a holidic food source for stable isotope labelling of amino acids in fruit flies. SILAF circumvents undesirable amino acid conversion by directly feeding Lys-6 to flies instead of pre-labelled yeast. We achieve virtually complete labelling already in second instar larvae, whereby the only efficiency limitation is the enrichment of the heavy aminoacid by the manufacturer. Sigma-Aldrich guarantees 99 atom % ^13^C, and we achieve 99.5% labelling with Lys-6.

We demonstrate in three biological applications that SILAF can be used for high-accuracy quantitative proteomics. Firstly, in agreement with previous work (7), by applying small-scale fractionation of a female/male fly mix we identified a number of differentially expressed proteins, including a prominent enrichment of yolk sack proteins 1, 2 and 3 in female flies, which have been shown to be exclusively present in the female fat body (15). Secondly, we applied a ratio-to-ratio analysis onto DmLRPPRC1 knockdown flies and performed single-shot proteomics. We found decreased protein levels in complex I, III and IV, but normal complex II and V. Interestingly, this corresponds to 7-week-old LRPPRC knockout mice (21) and is in agreement with biochemical complex activities performed in these fly lines before (16). Thirdly, high pH reversed-posed fractionation and phospho-peptide enrichment gave deep insights into enzymatic processes that were changed upon feeding the holidic diet compared to yeast medium. Using MaxQuant, we found 12,221 phosphorylation sites, whereof 7,496 were classified as high-confidence sites. The software could quantify 1,356 occupancies, which we used to get insight into metabolic pathways. Overlaying 4,749 quantified proteins in one or both replicates with phospho-occupancies revealed an enzymatic shift towards increased amino acid turnover, decreased glycolysis and fatty acid metabolism with increased gluconeogenesis and glycogen breakdown. The two latter mechanisms point towards a low dietary availability of a carbon source (28). However, increasing the sucrose to amino acid ratio correlated with decreased lifespan and egg-laying of flies (9). Yet, it appears that energy metabolism in these flies relies more on the pool of free amino acids, as both proteasomal and ribosomal machineries are increased.

Finally, we traced incorporation of the heavy Lys-6 label in both developing larvae and flies. Labelling in larvae happens fast and can be considered sufficient for accurate quantitative proteomics already at second instar stage, indicating rapid growth and protein turnover. This underlines that the fly is a valuable SILAC candidate. Cells for instance need two weeks of labelling time, mice two to four weeks with repeated changes of medium or chow, respectively. Larvae, in contrast are developing within two weeks in the same vial without the need to transfer them onto fresh food until hatching. Following Lys-6 incorporation, we find that the mitochondrial metabolic processes oxidative phosphorylation (OXPHOS) and tricarboxylic acid cycle incorporate the heavy label slower than the majority of cellular proteins, with only the ribosome being labelled even slower. In contrast, adult flies resemble a much more steady-state system in comparison to larvae, and thus show in general much slower labelling, in which ribosomal proteins turn over considerably faster than OXPHOS subunits. This underlines that turnover of functional protein categories correlates with the fly developmental stage.

### Further fields of application

We show that SILAF allows for concomitant fractionation of a two-sample peptide mix, enrichment of phospho-sites and subsequent mass-spectrometry to obtain a deep differential phospho-proteome. The method can be adapted to any post-translational modification (PTM) with enrichment methods in place, including epigenetic histone PTM profiling. Beyond that, SILAC has been used to decrease the false-positive rate of PTM screens like methylation, known as heavy-methyl SILAC (29). In this method, one population of cells is grown in the presence of methionine-4, of which the terminal methyl group in the methionine side chain is labelled. This methyl group is transferred via S-Adenosylmethionine onto a target by methyltransferases. Mixing a light and heavy fraction gives a 4 Dalton peptide mass shift only if a methyl-group truly exists. We think that the ease of SILAF allows this strategy to be applied on further amino acid-based PTMs like glutathionylation, ubiquitination or sumoylation.

SILAC affinity purification is a popular method to determine the interaction partners of a tagged bait protein (4, 30, 31) as it can be used to cancel out unspecific binding to antibodies and beads, thus decreasing noise and the number of false positive hits. *Drosophila* provides a simple and well-established model for transgenic expression of tagged proteins (32). With SILAF, co-immunoprecipitation studies can be performed in an *in vivo* model in a short timespan at reasonable costs.

As shown above, Lys-6 incorporation can be traced in time in both larvae and flies. A combination of pulse-chase setups with genetic defects can give insights into quantitative protein turnover rates, metabolic conversion routes of labels into proteinogenic amino acids and ultimately disease mechanism. Furthermore, we envision a spatial tracing by performing organ-specific mass spectrometry. The experimental setup can be enlarged for drug-protein interactions, bringing existing *Drosophila* drug screening strategies (33) together with preclinical pharmacoproteomics (34).

We describe an implementation of SILAF in its basic form and this can be a primer for adapting state-of-the art SILAC flavours in *Drosophila,* which have been already used in other model organisms. For instance, neutron encoded (NeuCode) SILAC allows sample multiplexing beyond the here presented two-sample mix (35, 36). The sample preparation requires complementation of growth media with amino acid isotopologues, which are commercially available. The implementation with SILAF holiday food in *Drosophila* is easy to envision.

### Limitations

The artificial food source causes delayed pupation of fly larvae, although it does not affect lifespan as shown before (10). That indicates that either the ratio of metabolite groups is imbalanced or that the quantity of one or more compounds is not adequately titrated. This highlights that the controls of SILAF proteomic experiments have to be well chosen in order to avoid secondary noise caused by comparing different developmental stages. Future improvements of the holidic food source might counteract this problem. Furthermore, flies grown on holidic food are smaller and lighter (9). Mixing an equal number of flies, whereof one partition is grown on a different diet, and processing those together would result in a systematic decrease of the heavy peptide fraction. In our approach, we decided to combine proteins only after reduction and alkylation based on protein mass, although this slightly reduces the power of SILAC in combining two experimental conditions very early on in sample preparation, thus increasing the likelihood of systematic errors (36).

We see 3.6% of all measured proteins significantly changed in flies grown on holidic diet compared to yeast, which suggests that the holidic diet does not cause a major caloric restriction or metabolic defect. We do observe, though, that the activation of alternative metabolic pathways can affect a mutant phenotype. For instance, DmLRPPRC1 knock-down larvae did not hatch on artificial food, whereas they do on yeast medium, although delayed (16). One option is to include climbing larvae in the analysis, although climbing is more difficult to stage than a hatching fly. A ratio of ratios analysis against a heavy standard, which can be easily generated and re-used, can help circumventing this problem, although abolishes one of the SILAC advantages of combining two or more samples in one run. Yet, if facility costs and time are not the limiting factor, the possibility to obtain a differential proteome or phospho-proteome upon fractionation is maintained, which guarantees the usage of SILAF for post-translational modification studies when a mutant is viable on at least one food source. Advancing SILAF further into NeuCode-SILAF would further mitigate this problem.

## MATERIAL AND METHODS

### Fly stocks and husbandry

Experiments were performed on a white Dahomey (wDah) fly line. Ubiquitous knockdown of bsf/DmLRPPRC1 was achieved by crossing the previously published fly line w;;UAS-bsfRNAi#1 (16) to a daughterless-GAL4 driver line. All stocks were backcrossed into the wDah genetic background for at least six generations. Parental flies were put to lay for 8 hours at an approximate density of 50 males to 50 females on either yeast or holidic medium. Experiments were performed on a mix of male/female stage three larvae or on one-day old male flies, if not indicated differently. Flies were maintained at 25°C and 60% humidity on a 12h:12h light/dark cycle.

### Holidic and yeast medium

Yeast medium was prepared with a standard yeast-sugar-agar composition. Holidic medium was prepared as described by Piper et al. (9, 10). Detailed recipes are available from the authors upon request. For SILAF labelling, heavy L-lysine-[^13^C[4/_6]_^15^N[0/_0_]] HCL (Sigma-Aldrich, 643459) and heavy L-arginine-[^13^C_6_,^15^N[0/_4_]] HCL (Sigma-Aldrich, 608033) were used. Medium was stored for up to 10 days at 4°C. All procedures were performed using semi-sterile techniques.

### Peptide sample preparation

Larvae were collected by either picking them with a sterile inoculation needle or by rinsing the food with distilled water. All larvae were cleaned in distilled water, soaked dry, collected in 1.5mL low-binding tubes (Eppendorf, 0030108116) and snap-frozen in liquid nitrogen. Flies were collected by a short carbon dioxide anaesthesia, collected in low-binding tubes and snap-frozen in liquid nitrogen. For all experiments with the exception of the phosphorylome, 10 insects were used as either a random mix of female/male larvae or as male flies only if not stated differently. SILAF samples were thus a mix of 5 light and 5 heavy insects. All samples were stored at −80°C before processing.

For protein extraction, samples were covered with 200μL 6M guanidine hydrochloride (GuCl, Sigma-Aldrich G3272) in 20mM Tris(2-carboxyethyl)phosphine hydrochloride pH 8.0. All samples were homogenised with a Teflon-coated pestle for about 10 seconds. Samples were incubated at room temperature (RT) for 10 minutes, then sonicated at a 10 seconds on/off cycle in ice-cold water four times, followed by a further incubation of 10 minutes at room temperature. A centrifugation step for 5 minutes at maximum speed pelleted most of the insoluble chitin, although a floating layer can occur. 180μL of supernatant without any visible fly chitin was transferred to a new 1.5mL low-binding tube. An aliquoted 1:10 dilution in water was quantified with Pierce BCA protein assay (ThermoFisher Scientific Cat. No. 23225). While the protein assay was incubating at 37°C in the dark, proteins were first reduced for 30 minutes at 55°C with 5mM dithiothreitol (DTT), cooled down briefly on ice and then alkylated for 15 minutes at room temperature in the dark with 15mM 2-chloroacetamide (37). 50μg of each light and heavy protein sample was then mixed, transferred to a new 2mL low-binding tube, and brought to 1.8mL with 110mM Tris pH 8.5. Label-free samples were used at 100μg. Either Pierce lysyl endopeptidase LysC, MS grade (ThermoFisher Scientific Cat. No. 90307) or Pierce trypsin, MS grade (ThermoFisher Scientific Cat. No. 90057) was added at 1:50, equivalent to 2μg of protease and incubated over night for a maximum of 18 hours lying at 37°C and mild shaking.

Samples were acidified with 1.2% formic acid, spun at 3,000g and room temperature for 10 minutes and desalted on 3mL Empore™ SPE cartridges (Sigma-Aldrich 66872-U), following the manufacturers recommendations. In brief, the column was conditioned with 200μL methanol and equilibrated with 500μL water. The sample was loaded followed by 1mL acidified sample buffer and two washes of 500μL each, the first one with water, the second one with 0.5% formic acid. The peptides were eluted off the column with 300μL of 40% acetonitrile two times and lyophilised in a SpeedVac (Savant SC110A vacuum concentrator with refrigerated vacuum trap). Peptides were resolubilised in 20μL of 0.5% formic acid and quantified on a NanoDrop 1000 spectrophotometer at 280nm. Following this protocol, anticipated amounts are between 20 – 35μg of peptides upon 100μg protein input. Peptides were dried again, stored at −80°C and sent for mass spectrometry analysis on dry ice.

For the phosphorylome, 250 male flies were collected for each fraction and replicate. Phosphorylome flies were split into two equal portions, 860mg GuCl was added and brought to 1.5mL with 20mM Tris pH 8.0 to obtain a final concentration of 6M GuCl. Homogenisation was performed as described above and 1.3mL were transferred to a 15mL tube. Reduction and alkylation were performed as described above and samples were diluted 1:10, which obtained about 1mg/mL. Ten milligram of protein extract from each sample was combined and subjected to digestion by 40μg endoproteinase LysC at 37°C for 12 hours. Next, 10μg of LysC was added and the digestion was carried for four more hours. Peptide samples were acidified using formic acid to a final concentration of 1% and centrifuged at 5000 rpm for ten minutes. The supernatant, containing peptides, was desalted by solid phase extraction using C18 Sep-Pak cartridge, 200 mg (Waters, WAT054945). The cartridge was conditioned with 1mL methanol, 1mL 60% acetonitrile 1% formic acid, and twice with 1mL 1% formic acid. Next sample was applied and washed twice with 2mL 1% formic acid. Finally the peptides were eluted using 600μL 60% acetonitrile 1% formic acid dried in a vacuum centrifuge. For large pH reversed phase separation, the peptide pellet was dissolved in 50% ACN, 0.01M using 10 min sonication in a water bath. The solution was diluted to 5% ACN, 0.01M ammonium bicarbonate, centrifuged for 10 min at 20,000 rcf on a tabletop centrifuge and 95% of the solution was separated by large-scale high pH reversed phase separation.

### Small-scale high pH reversed phase separation

Twenty micrograms of peptides derived from the SILAF comparison of male and female fly were separated on a 150mm, 300μm, 2μm C18, Acclaim PepMap (Product No. 164537, Thermo Fisher Scientific) column using a Ultimate 3000 (Thermo Fisher Scientific). The column was maintained at 30°C. Buffer A was 5% acetonitrile 0.01M ammonium bicarbonate, buffer B was 80% acetonitrile 0. 01M ammonium bicarbonate. Separation was performed using a segmented gradient from 1% to 50% buffer B, for 85min and 50% to 95% for 20 min with a flow of 4μL. Fractions were collected every 150 sec and combined into nine fractions by pooling every ninth fraction. Pooled fractions were dried in Concentrator plus (Eppendorf), resuspended in 5 μL 0.1% formic acid from which 2 μL analysed by LC-MS/MS.

### Large-scale high pH reversed phase separation and phosphopeptide enrichment

Peptides derived from each biological replicate were separated on a 4.6 x 250 mm ZORBAX 300 Extend-C18, 5μm, column (Agilent Technologies) at a 1 ml flow rate using a NGC Quest 10 chromatography system (Bio-Rad). Buffer A was 5% acetonitrile 0.01M ammonium bicarbonate, buffer B was 80% acetonitrile 0.01M ammonium bicarbonate. Buffers were prepared with LC-MS grade water. Peptide separation was performed using a segmented gradient from 5% to 27% buffer B for 65 min and from 27% to 45% buffer B for 30 min. Eluting peptides were collected for 72 min using a BioFrac fraction collector (Bio-Rad). Fraction collection pattern was set to row and fraction collection size was set to 0.75ml. In total, 96 fractions were collected in a 1.2ml V-shaped 96 well plate (Biotix). Upon sample collection, plates were dried overnight in Concentrator plus (Eppendorf) and peptides were resuspended in 20 μL 80% acetonitrile, 2% trifluoroacetic acid. All 96 fractions were concatenated into 12 fractions by combining every 12^th^ fraction. One-tenth of the volume was dried, resuspended in 5 μL 0.1% formic acid from which 2 μL analysed by LC-MS/MS. The remaining solution was diluted with lactic acid (GL Sciences) and phosphopeptides were enriched with 3mg TiO2 tips (GL Sciences) using the manufacturer’s instructions. Enriched phosphopeptides samples were dried and resuspended in 5 μL 0.1% formic acid from which 2 μL were used for analysis by LC-MS/MS.

### LC-MS/MS analysis

Peptides were separated on a 25 cm, 75 μm internal diameter PicoFrit analytical column (New Objective) packed with 1.9 μm ReproSil-Pur media (Dr. Maisch) using an EASY-nLC 1200 (Thermo Fisher Scientific). The column was maintained at 50°C. Buffer A and B were 0.1% formic acid in water and 80% acetonitrile, 0.1% formic acid, respectively. Peptides were separated on a segmented gradient from 6% to 31% buffer B for 82 min, and from 31% to 44% buffer B for 5 min at 200nl / min. Eluting peptides were analysed on a QExactive HF mass spectrometer (Thermo Fisher Scientific). Peptide precursor mass to charge ratio (m/z) measurements (MS1) were carried out at 120000 resolution in the 350 to 1500 m/z range. The top seven most intense precursors, with charge state from two to six only, were selected for HCD fragmentation using 27% normalised collision energy. The m/z of the peptide fragments (MS2) were measured at 30000 resolution, in profile mode, using an AGC target of 2e5 and 80 ms maximum injection time. Upon fragmentation precursors were put on an exclusion list for 45 sec.

Phosphopeptide enriched samples were analysed using two technical replicates. For the first replicate, peptides were separated with a gradient from 6% to 31% buffer B for 70 min, and from 31% to 44% buffer B for 17 min, using MS parameters as described above. For the second replicate, peptides were separated with a gradient from 6% to 31% buffer B for 53 min, and from 31% to 44% buffer B for 4 min. The MS parameters were as described, except that the MS2 of the top five most intense precursors were measured at 60000 resolution with 110 ms injection time.

Peptide fractions from the small-scale high pH reversed phase separation were analysed on an Orbitrap Tribrid Fusion (Thermo Fisher Scientific) with the same MS parameters as described above, except for centroid mode of acquisition in MS2.

### Quantification and statistical analysis

Raw spectra were mapped in batches for either female/male comparison, the holidic phosphorylome and proteome, quality control and DmLRPPRC1 or proline-6 conversion with the free software MaxQuant version 1.6.1.0 (38) against the canonical and isoform sequences of the fruit fly proteome (UP000000803_7227 from UniProt, downloaded in April 2016). The reference database was complemented by MaxQuant’s own default contaminant library. Methionine oxidation and N-terminal acetylation were set as variable modifications. *In silico* digestion was performed with LysC/P as a protease. A false discovery rate of 0.01 was chosen for peptide spectra matches. Label-free samples were quantified with default LFQ settings.

Quality control of the mapped proteinGroup.txt file was performed separately for each study in Perseus version 1.5.0.0 (39). Normalised H/L ratios or label-free values were imported for each sample and proteins marked as “reverse”, “only identified by site”, and “potential contaminant” were excluded. Incorporation rates were calculated from not-normalised intensities by: Intensity Heavy / (Intensity Heavy + Intensity Light). Fold changes between two fractions were derived from normalised Heavy / Light ratios. For the DmLRPPRC1 ratio/ratio analysis, the following quotient was calculated: Normalised ratio of (Heavy standard / Light wDah control) / Normalised ratio of (Heavy standard / Light DmLRPPRC1-knock down)) = Normalised bfs-knock down / wDah control with subsequent log2 transformation. All other SILAF values were log2-transformed before mean values were calculated. Occupancies of phosphorylation sites per replicate were calculated as occupancy Heavy / occupancy Light and are presented as log2-transformed values. LFQ values were imported and mean values were calculated on not transformed data. For better visualisation, LFQ intensities were log10 transformed.

The proteinGroup.txt file and the Perseus output were imported into R version 3.4.2 running in a RStudio environment, version 1.1.383. The R code is publicly available via GitHub: https://github.com/Zinksulfid/SILAF_public.git. Figures were plotted with ggplot2 version 2.2.1.9000 and extension packages, see the original script. Adjusted p-values for plotting were calculated as FDR with limma moderated t-test, which is part of the limma package version 3.32.10.

Gene Set Enrichment analyses and overrepresentation analyses were performed with the WebGestalt 2017 online application (40) by uploading a ranked file containing Entrez Gene identifiers and mean log2-fold changes. Uniprot IDs were converted to Entrez Gene IDs using the package UniProt.ws version 2.16.0 in R. Mouse orthologues of fly OXPHOS subunits were identified with DIOPT – DRSC Integrative Ortholog Prediction Tool (41).

A description of statistical parameters used in each experiment is provided in the figure legends.

### Data and software availability

The mass spectrometry raw files and a summary of the data analysis is available from the authors upon request.

## ACKNOWLEDGEMENTS

The authors would like to thank Thomas Colby (MPI Biology of Aging) for developing the large and small-scale high pH reversed phase separations and Matthew Piper (Monash University) for advice on the holidic food. This study was supported by the Swedish Research Council [AW (VR2016-02179), AWe (VR2016-01082)], the Knut & Alice Wallenberg Foundation [AW and AWe (KAW 2013.0026)), the European Research Council [AW] and the Max Planck Society [IA]. FAS is part of the mentor program of the Studienstiftung des deutschen Volkes. AWr is a Ragnar Söderberg fellow of medicine.

## AUTHOR CONTRIBUTIONS

FAS, IA, CF, and AWr had the idea, conceptualized the study and planned experiments. FAS and DM performed fly work. FAS and IA prepared peptides, IA performed fractionation and mass spectrometry. FAS and IA analysed the data and wrote R code. CF, AWr, FAS, IA and DM conceived and wrote the manuscript. AWe contributed with reagents. IA, AWr and CF supervised all aspects of the study. All authors reviewed and edited on the manuscript before submission.

## FIGURE LEGENDS

**Figure S1:**
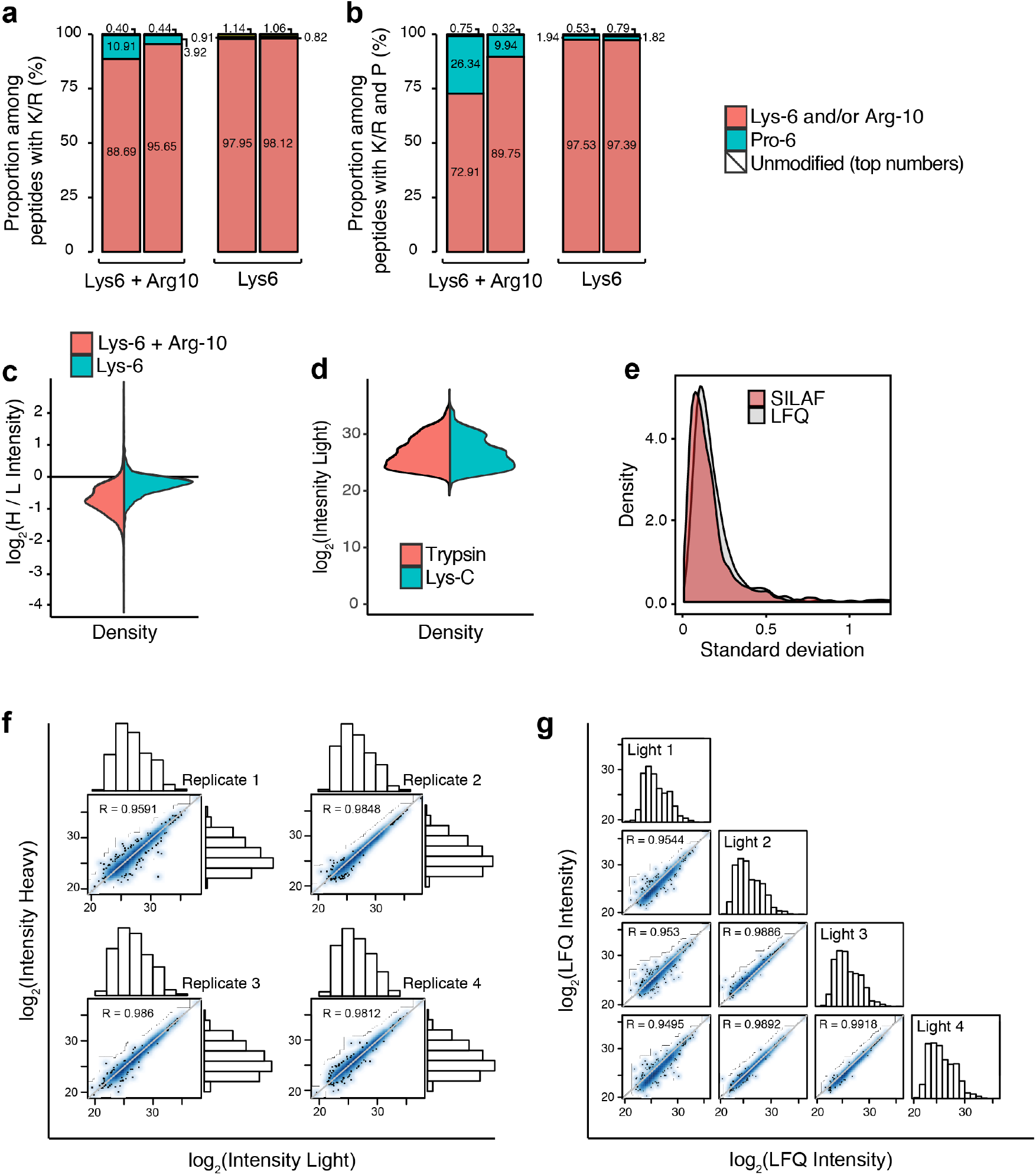
Quality control of the SILAF method. (a) Proportion of labelled peptides in flies (n = 2) grown on medium with either Arg-10 and Lys-6 (left) or Lys-6 (right), proportion of unlabelled and Pro-6 containing peptides among peptides containing lysine or arginine. (b) Data as in (a) for all peptides that contain proline. (c) The Arg-10 label causes a partial loss in intensity of identified peptides in comparison to Lys-6. Light (L) peptides were derived from wDah grown on holidic diet (n=3). The heavy (H) fraction was labelled with either Lys-6 and Arg-10 and digested with trypsin, or Lys-6 only and digested with Lys-C. The graph is based on not normalised intensity values. (d) Lys-C peptides have similar abundance profile to tryptic peptides. The plot shows the not normalised intensity of light SILAF fractions derived from light/heavy protein mixes from holidic food, digested either with trypsin or Lys-C (n=3). Samples as of (c). (e) Standard deviation per quantified protein of normalised H/L ratios of a holidic diet proteome (SILAF) and label-free quantification intensities of the light fraction of the same sample (LFQ; n=4 per density plot). (f) Corresponding scatter plots to (e) for SILAF and (g) LFQ quantifications with respective Pearson’s correlation coefficients.

**Figure S2:**
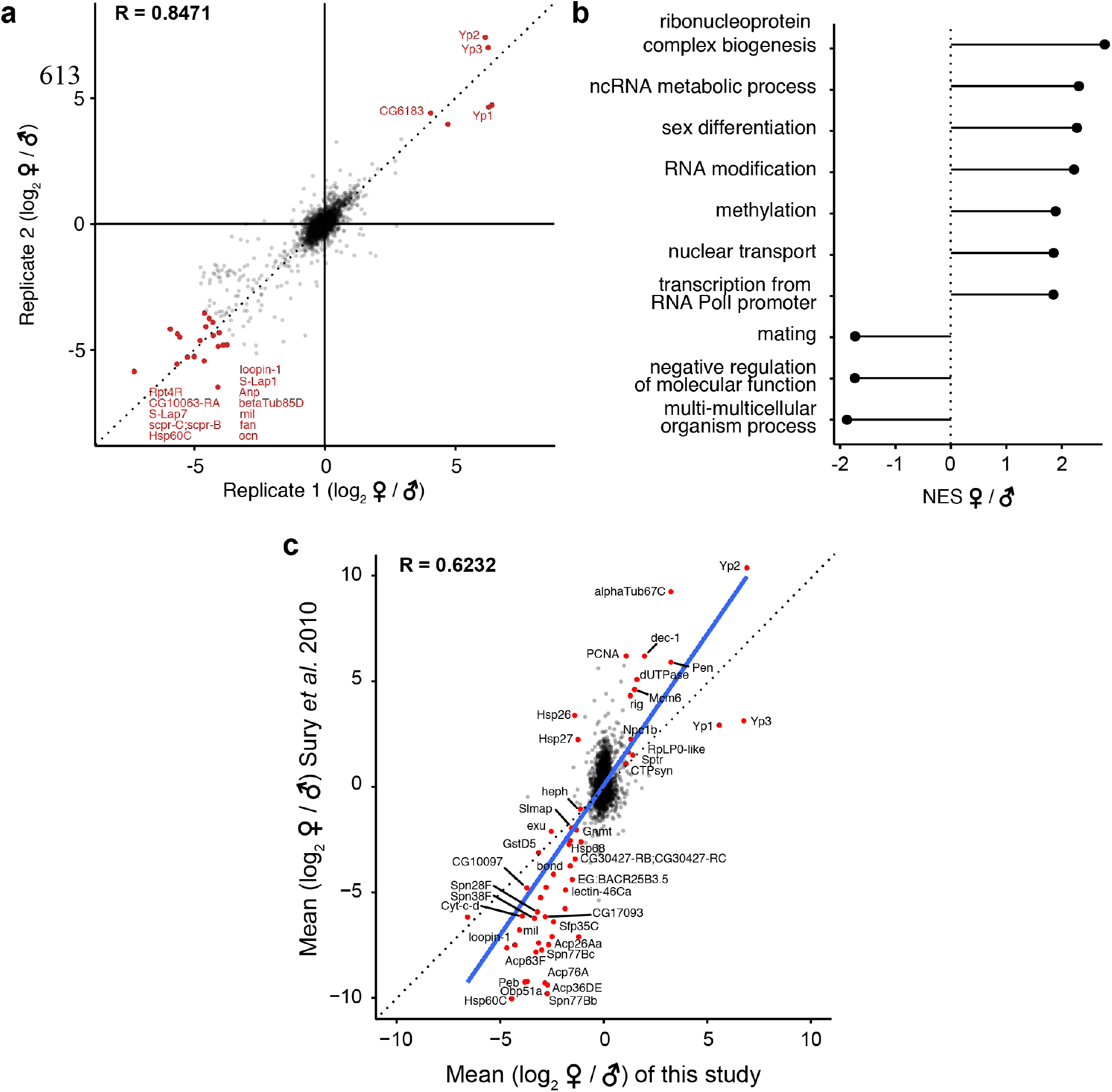
SILAF based comparison of female and male fly proteomes. (a) Overlap of two biological SILAF replicates. Replicate 2 was performed with inverted labels. Red dots show targets with an absolute log2-fold change of more than 4 in both replicates. (b) Gene Set Enrichment Analysis of all proteins in (a) and plotted with their normalised enrichment score (NES). (c) Comparison of the results of this study obtained with SILAF to previous data that was generated from flies grown on labelled yeast (Sury et al., 2010). A linear trend line in blue. R is Pearson’s correlation coefficient.

**Figure S3:**
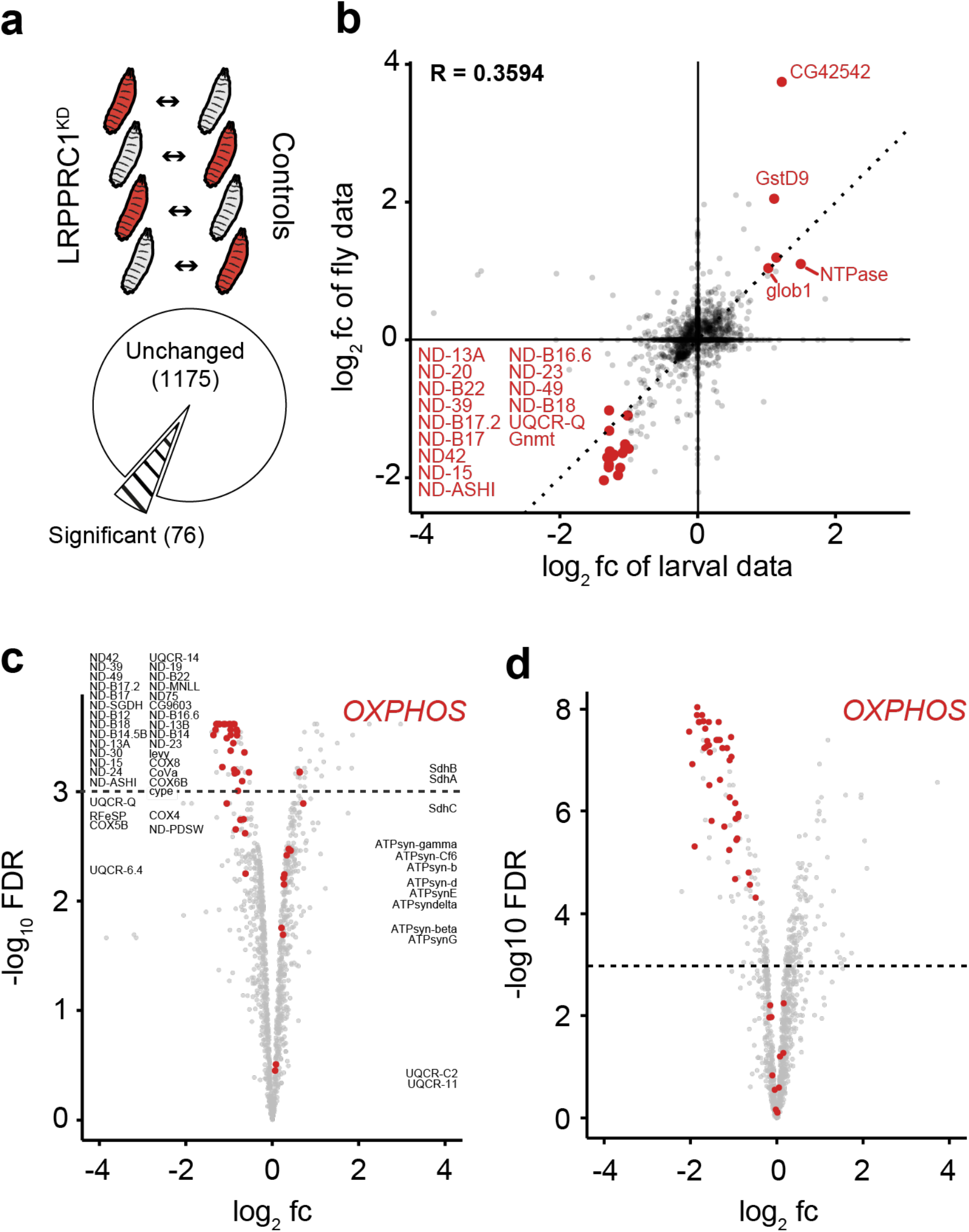
SILAF for accurate proteome quantification used in a DmLRPPRC1 knock-down model. (a) Classical SILAF based on direct labelling of biological samples and label swap. Protein extracts from light and heavy larvae (n=4) were mixed, analysed by single-shot LC-MS/MS followed by a one-sample moderate t-test. (b) Comparison of fold changes derived from a classical approach (larval data, n= 4) and the spike-in SILAF approach (fly data, n = 4 for each controls and knock-down samples). R is Pearson’s correlation coefficient. (c) OXPHOS members (red) in a volcano plot of the classical SILAF dataset with gene names in the proximity of the data point. (d) OXPHOS members (red) in a volcano plot of the results from the SILAF spike-in experiment.

**Figure S4:**
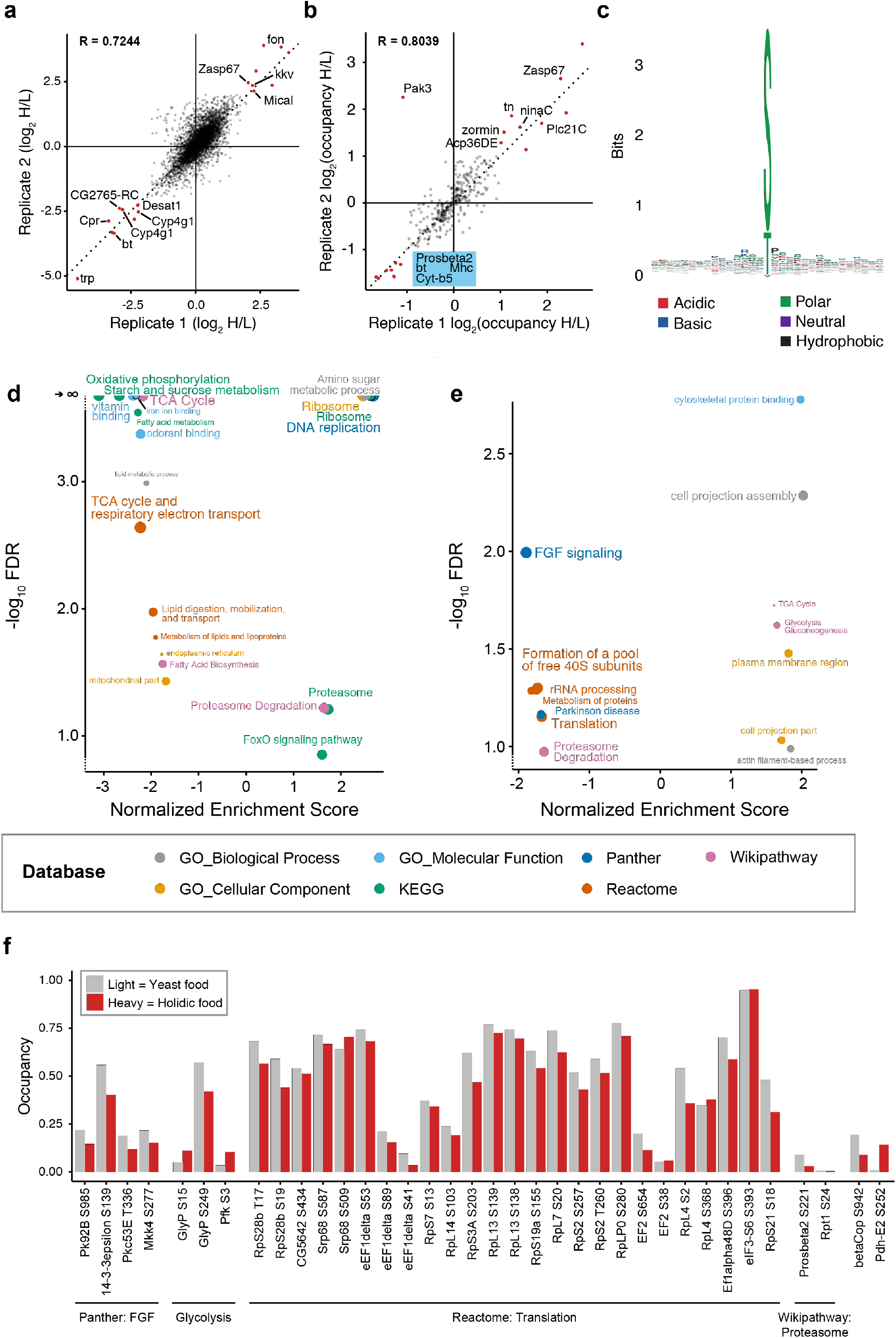
Differential phosphorylome and proteome of holidic versus yeast food-grown flies. (a) Overlap of heavy (H, holidic diet) to light (L, yeast diet) ratios of phospho-peptides in two replicates. (b) Differences in phospho-occupancies in both replicates. R is Pearson’s correlation coefficient. (c) Phospho-site enrichment (n = 7496). (d) Gene Set Enrichment Analysis of proteins shown in Fig. 4b (n = 2) and (e) changes in the phosphorylation occupancy of proteins shown in Fig. 4c against several databases as indicated (n = 2). Cut circles have an FDR < 0.1 %. (f) Occupancies of selected targets as identified in Fig. 4e and Supplementary Fig. 4b. Mean values of two replicates.

